# Network Toxicology, Molecular Docking, and Molecular Dynamics Simulations Reveal the Mechanism of Tetrabromobisphenol A in Bullous Pemphigoid

**DOI:** 10.64898/2026.03.27.714718

**Authors:** Kuo Sun, Yuxin Liu, Zhao Hanqing

## Abstract

Bullous pemphigoid (BP) is an autoimmune blistering disease with a growing incidence, and environmental factors are receiving increasing attention. Tetrabromobisphenol A (TBBPA), a widely used brominated flame retardant, is a significant environmental pollutant. However, the molecular mechanisms by which TBBPA contributes to BP pathogenesis remain unclear. This study integrated network toxicology, molecular docking, and molecular dynamics (MD) simulations to systematically investigate the molecular mechanisms of TBBPA-induced BP. Using network toxicology, we identified 797 potential targets of TBBPA and 446 BP-related targets. A Venn diagram analysis revealed 48 common targets. Protein-protein interaction (PPI) network and topological analyses further identified five core hub targets: TNF, CXCL8, MMP9, ICAM1, and ITGB1. Gene enrichment analysis indicated that these targets were significantly enriched in immune-inflammatory pathways, such as leukocyte migration, inflammatory responses, and the IL-17 signaling pathway, as well as in various pathogen infection and cancer-related pathways. Molecular docking revealed that TBBPA stably binds to all five core targets with binding energies ≤ -5 kcal/mol, driven primarily by hydrophobic interactions and π-π stacking. Subsequent MD simulations confirmed that TBBPA complexes with TNF, CXCL8, and MMP9 remained stable throughout the 100 ns simulation. The overall protein structures remained compact, and the ligands were effectively encapsulated within the binding pockets, forming stable networks of hydrogen bonds and hydrophobic interactions. In conclusion, this study, for the first time, proposes a systematic molecular framework using integrated computational biology. Our findings suggest that the environmental pollutant TBBPA may act as a potential risk factor in BP pathogenesis by targeting core proteins (TNF, CXCL8, and MMP9). These interactions potentially disrupt critical signaling pathways related to immune inflammation, cell migration, and tissue remodeling. This study offers a novel mechanistic hypothesis regarding environmental chemical exposure in autoimmune blistering diseases, although further experimental validation is required.

**Highlights:**

- Network toxicology identified 48 common targets linking Tetrabromobisphenol A(TBBPA) exposure to Bullous Pemphigoid (BP).
- Five core targets (TNF, CXCL8, MMP9, ICAM1, ITGB1) were screened as potential mediators.
- TBBPA stably binds to TNF, CXCL8, and MMP9 with binding energies ≤ -5 kcal/mol.
- Molecular dynamics simulations confirm stable binding and structural integrity of complexes.
- This study provides a mechanistic framework for TBBPA as an environmental risk factor in BP.

## 1. Introduction

Bullous pemphigoid (BP) is a common autoimmune subepidermal blistering disease predominantly affecting the elderly. Its pathogenesis is complex, involving genetic, immune, and environmental factors. Individuals carrying genetic susceptibility alleles regulating immune function may have a significantly increased risk of BP upon exposure to specific environmental triggers [1].

Among various environmental factors, chemical pollutants have attracted considerable attention. Tetrabromobisphenol A (TBBPA) is a major brominated flame retardant widely used in products such as electronic equipment, making it a ubiquitous environmental pollutant [2]. Humans can be exposed through various routes, including dermal contact. We hypothesize that exposure to TBBPA may represent an environmental risk factor for BP by interfering with specific molecular pathways; however, these molecular regulatory mechanisms have not yet been elucidated.

Network toxicology, an extension of network pharmacology, enables systematic screening for common targets between environmental chemicals and diseases and facilitates the construction of protein-protein interaction (PPI) and pathway networks [3]. To validate predictions at the molecular level, this study employed molecular docking to construct binding conformations of TBBPA with core candidate targets [4]. Molecular dynamics (MD) simulations were also conducted to analyze the stability and dynamic interactions of these complexes under simulated physiological conditions, revealing binding affinity and interaction mechanisms at the atomic level [5].

This study integrated network toxicology, molecular docking, and MD simulations to investigate the potential molecular mechanisms by which TBBPA may contribute to the pathogenesis of BP. It screened key targets, identified relevant signaling pathways, and characterized stable binding interactions, thereby providing a theoretical basis for understanding environmental triggers of autoimmune blistering diseases.

## 2. Methods

### 2.1. Collection of TBBPA Targets

The structural information of TBBPA was obtained from the PubChem database. The structure was then submitted to the ChEMBL, TargetNet, and SwissTargetPrediction databases, with the species restricted to *Homo sapiens* [6-8]. Potential targets of TBBPA were subsequently retrieved. The results were merged and deduplicated, and target names were standardized using the UniProt database [9]. These targets were then used to construct the TBBPA target library.

### 2.2. Screening of BP-Related Targets

We searched for related targets using the keyword “BP” in the GeneCards, OMIM, and TTD databases [10-12]. Venn diagram analysis was then used to identify common targets between TBBPA and BP. The intersection was defined as the potential targets for TBBPA-induced BP.

### 2.3. PPI Analysis and Core Target Screening

The intersecting genes were input into the STRING (version 12.0) database, with the species restricted to *Homo sapiens* and the minimum interaction score set to high confidence (> 0.4). The results were imported into Cytoscape 3.10 to calculate network parameters and construct the PPI network [13]. Core targets were screened using the following criteria: degree > 2 × median, closeness centrality > median, betweenness centrality > median, and average shortest path length > median.

### 2.4. Gene Function and Pathway Enrichment Analysis of Target Proteins

To investigate the biological functions of potential targets in TBBPA-induced BP, Gene Ontology (GO) and Kyoto Encyclopedia of Genes and Genomes (KEGG) enrichment analyses were performed using R (version 4.4.0) and the clusterProfiler package (version 4.4.4) [14]. Gene symbols were converted to Entrez IDs using the org.Hs.eg.db database (version 3.15.0) [15]. GO analysis included biological process (BP), cellular component (CC), and molecular function (MF). KEGG enrichment was conducted using the enrichKEGG function with organism = “hsa”, based on the latest KEGG database [16]. Significance thresholds were set at p < 0.05 and adjusted p (FDR) < 0.05. Visualization was performed using clusterProfiler and ggplot2 (version 3.4.0) [17] to generate bar and bubble plots. KEGG results were presented as lollipop plots using ggpubr (version 0.5.0) [18]. All figures were exported in PDF format. These analyses systematically revealed key biological processes and signaling pathways associated with TBBPA-induced BP.

### 2.5. Molecular Docking of TBBPA with BP Core Targets

This study employed molecular docking to investigate potential binding modes between TBBPA and BP-related target proteins. Five core targets were selected: TNF, CXCL8, MMP9, ICAM1, and ITGB1. Docking simulations between TBBPA and each target were performed using AutoDock software to predict optimal binding conformations, binding energies, and interactions. Following docking, PyMOL software was used to visualize and preliminarily analyze the 3D structures of the complexes, confirming binding sites. Two-dimensional interaction diagrams were generated using specialized molecular analysis tools to systematically identify specific interactions (e.g., hydrogen bonds, van der Waals forces, π-π stacking) and amino acid residues involved.

### 2.6. MD Simulation of TBBPA with BP Core Targets

MD simulations were performed to further investigate the binding modes and dynamic stability of TBBPA with core targets. Based on docking results, the most stable complexes of TBBPA with three target proteins (CXCL8, MMP9, and TNF) were selected as initial structures.

All simulations were performed using the GROMACS 2023.2 software package and the CHARMM36 force field. The GROMACS 2023.2 package provides comprehensive tools for building, energy minimization, equilibration, and simulating the dynamic behavior of molecular systems [19]. Topology parameters for TBBPA were generated using the CGenFF program. Each complex was placed in a cubic water box containing TIP3P water molecules, and appropriate ions were added to neutralize the system. Each system was energy-minimized using the steepest descent and conjugate gradient methods. Systems were then equilibrated under NVT and NPT ensembles, each lasting 1 ns, at 300 K and 1 bar. Finally, an unrestrained MD simulation was conducted for 100 ns, using a 2 fs time step. Trajectory analyses included calculations of root mean square deviation (RMSD) of the complex and ligand, root mean square fluctuation (RMSF) of the protein, radius of gyration (Rg), number of hydrogen bonds between ligand and protein, and solvent-accessible surface area (SASA) of the ligand. All analyses were performed on the equilibrated portion of the trajectories.

## 3. Results

### 3.1. Identification of Targets for TBBPA-Induced BP

This study employed network pharmacology to systematically screen and identify potential targets of TBBPA-induced BP. Potential TBBPA targets were retrieved from three authoritative databases, TargetNet, ChEMBL, and SwissTargetPrediction, resulting in 797 unique targets after merging and deduplication. Concurrently, disease targets relevant to BP were identified using the keyword “BP” in three databases: OMIM, GeneCards, and TTD. After deduplication and verification, 446 BP-associated targets were obtained.

A Venn diagram intersection analysis was performed to identify core targets mediating TBBPA-induced BP. The combined dataset contained 1195 non-redundant targets, of which 749 targets (62.7%) were unique to TBBPA, 398 targets (33.3%) unique to BP, and 48 common intersecting targets (4.0%). These 48 intersecting targets were identified as potential core targets for TBBPA-induced BP (Fig. 1).

**Fig. 1.**
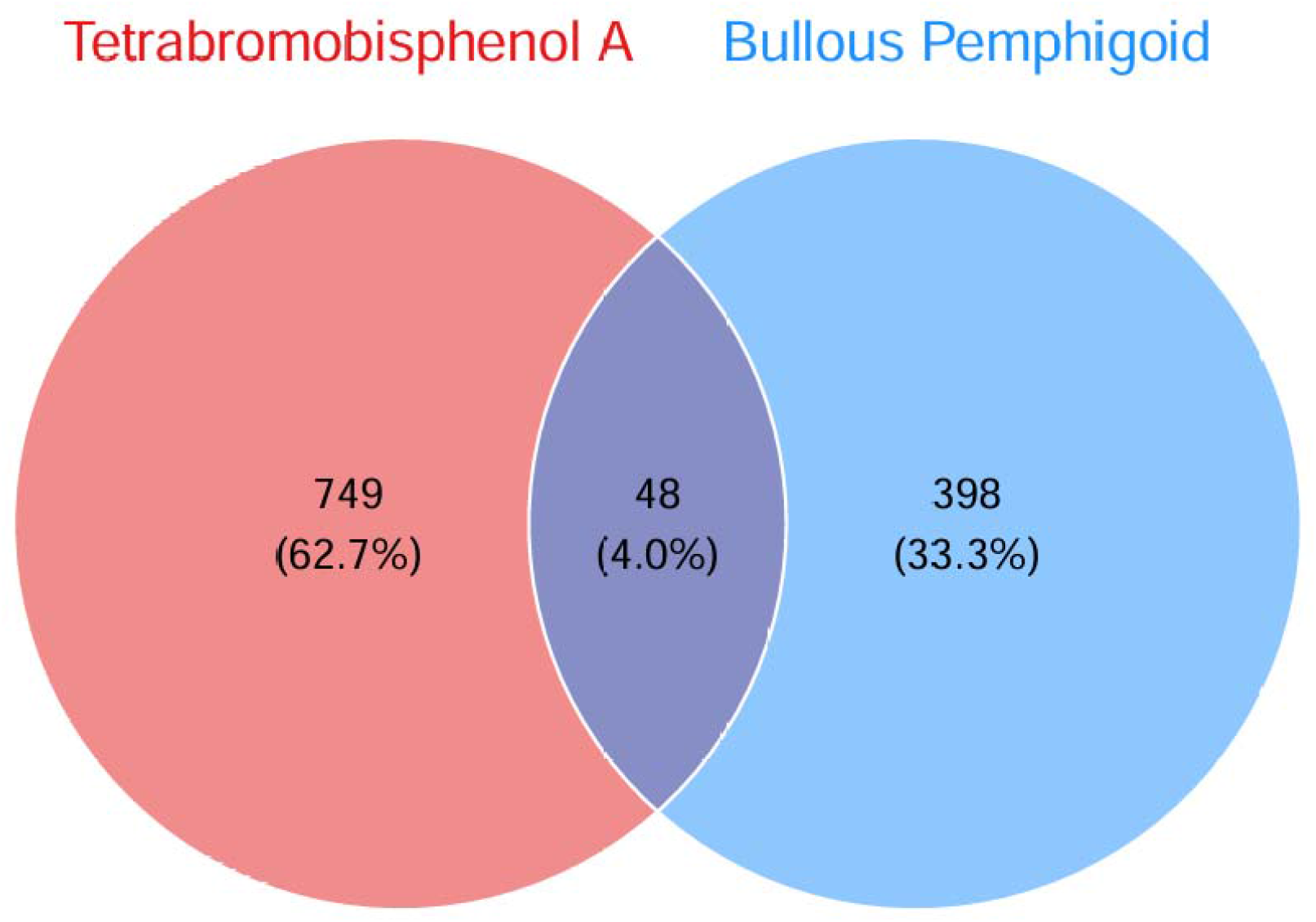
Venn diagram showing common targets of TBBPA and BP.

### 3.2. Construction of PPI Network and Screening of Core Regulatory Genes

The 48 potential targets for TBBPA-induced BP were imported into the STRING 12.0 database for PPI analysis. After removing isolated nodes, the resulting PPI network consisted of 46 nodes and 314 interaction edges, with an average node degree of 13.1. Cytoscape 3.10 software was used to analyze topological properties, including degree centrality, closeness centrality, and radiality, and visualize the network (Fig. 2). The top five core targets by degree centrality were TNF, CXCL8, MMP9, ICAM1, and ITGB1. Node size and color intensity positively correlated with degree centrality, reflecting their central roles in the network. Existing research has widely confirmed that proteins encoded by these five core genes critically regulate BP pathological processes, including innate and adaptive immune responses, inflammatory cascade initiation and amplification, neutrophil and lymphocyte chemotaxis and infiltration, extracellular matrix degradation, and dermal-epidermal junction integrity.

**Fig. 2.**
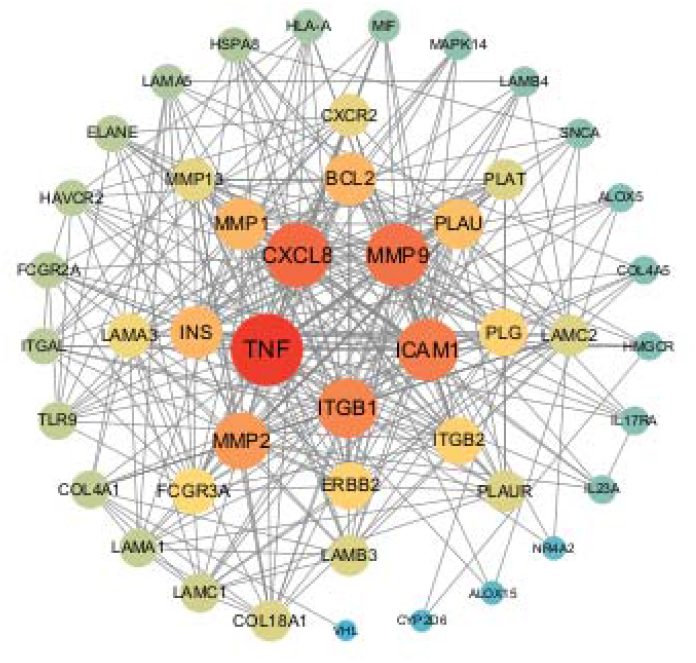
PPI network of core targets.

### 3.3. Target Function and Pathway Enrichment Analysis

#### 3.3.1. GO and KEGG Analysis of Targets

GO functional enrichment analysis was performed for the 48 potential targets associated with TBBPA-induced BP using R software (version 4.4.0) and the clusterProfiler package (version 4.4.4). Gene symbols were converted to Entrez IDs using the org.Hs.eg.db database (version 3.15.0). Applying significance thresholds (P < 0.05, adjusted P-value (FDR) < 0.05), the analysis identified 839 statistically significant GO terms, including 779 BP terms, 32 CC terms, and 28 MF terms. GO terms were sorted by ascending FDR values. The top 10 terms from each GO category (BP, CC, MF) were visualized using enrichment bar charts and bubble plots (Fig. 3).

**Fig. 3.**
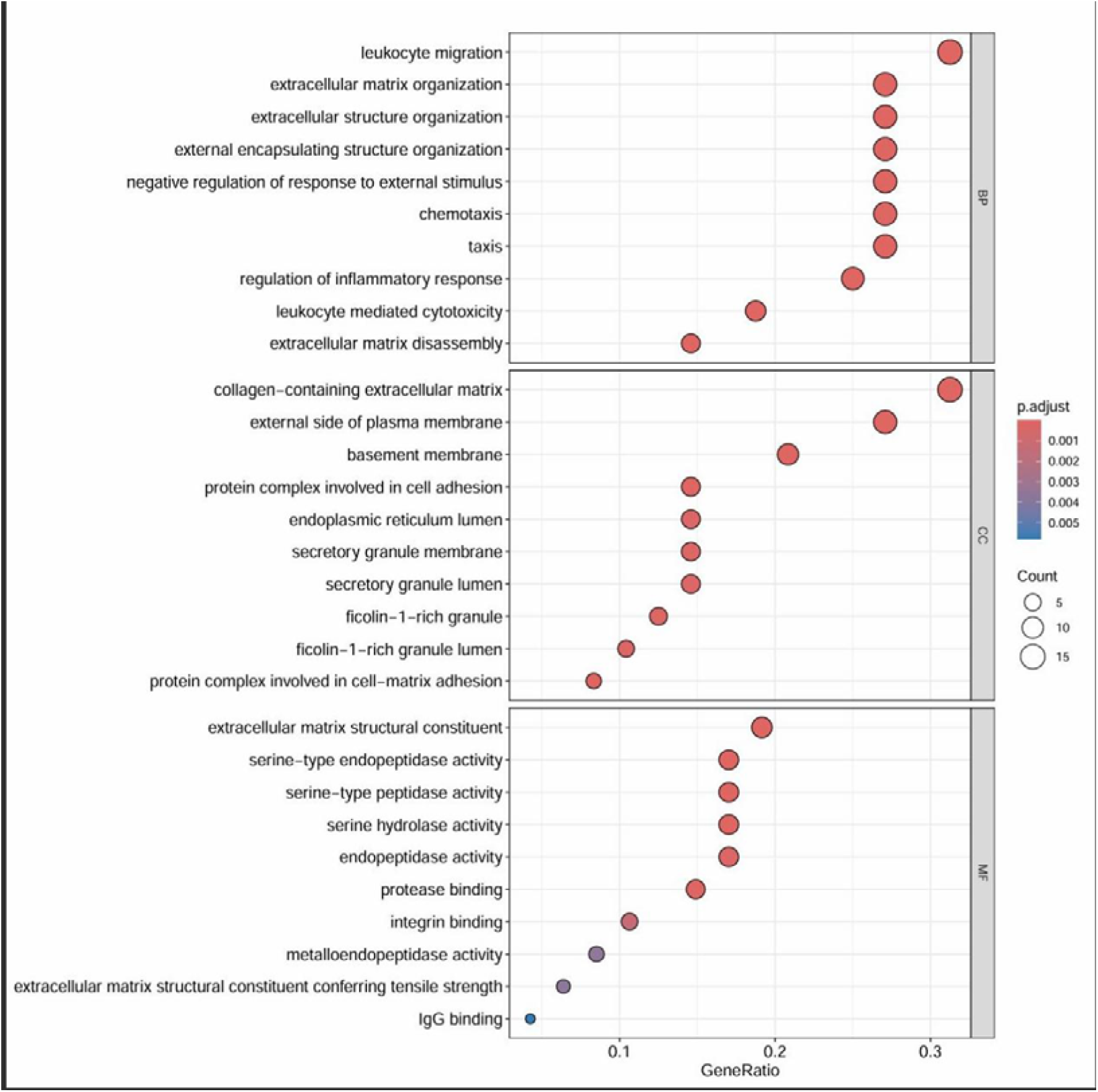
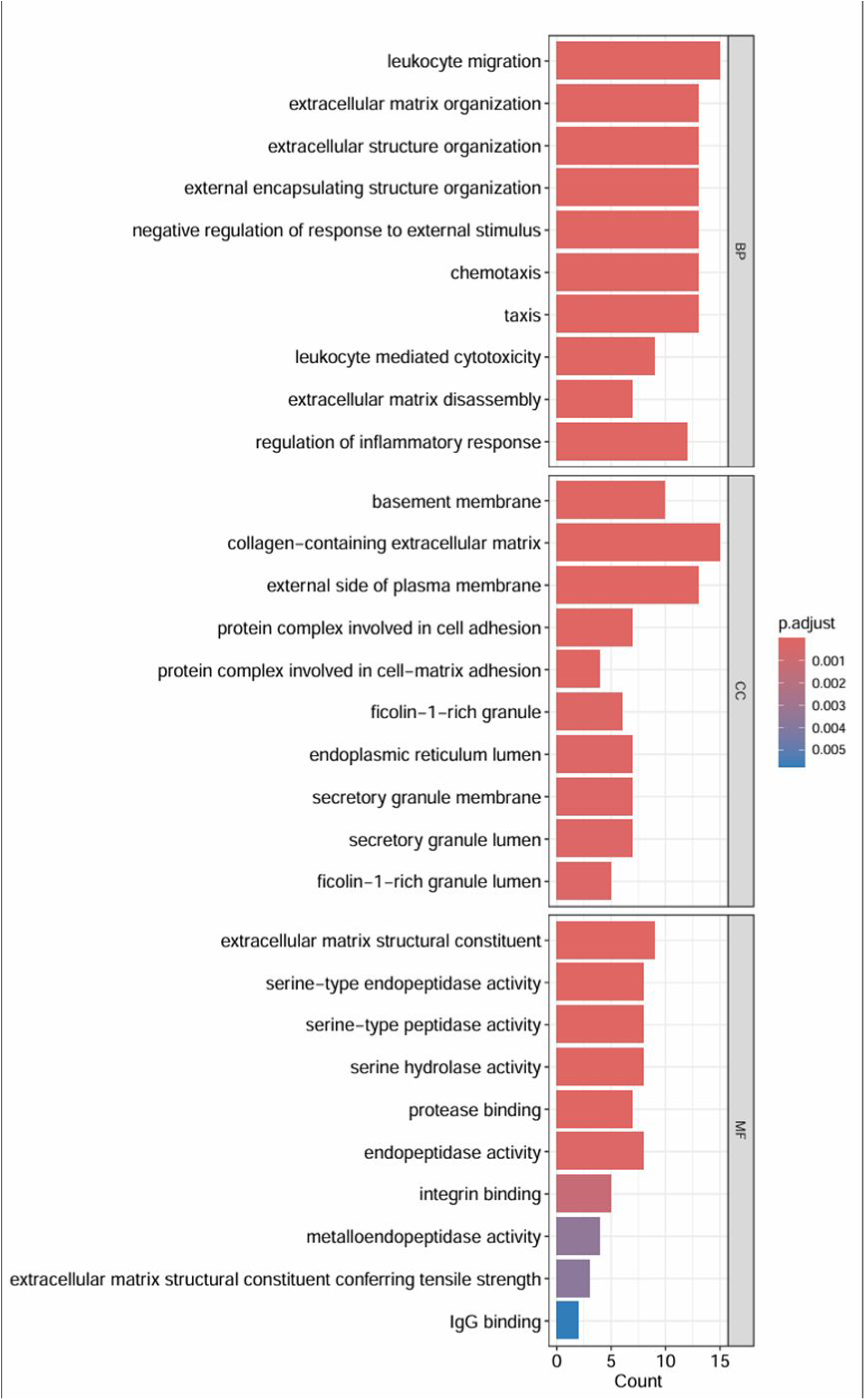
To elucidate the functional enrichment of genes intersecting between TBBPA and BP, GO analysis was performed, categorizing genes into BP, CC, and MF. Significant enrichment was identified for BP terms, including leukocyte migration and extracellular matrix organization; CC terms, such as basement membrane and collagen-containing extracellular matrix; and MF terms, particularly extracellular matrix structural constituent and serine-type endopeptidase activity. Low adjusted p-values (p.adjust) indicate statistical significance, while Count values represent gene numbers. These findings suggest that intersecting genes may participate in inflammatory processes and extracellular matrix remodeling, potentially related to BP pathology.

Additionally, KEGG pathway enrichment analysis was conducted for the 48 potential targets using the enrichKEGG function of the clusterProfiler package in R, based on the latest KEGG human database (organism = “hsa”). Using an FDR < 0.05 threshold, significantly enriched pathways were identified (specific numbers temporarily unavailable). Visualization of the top 30 signaling pathways with the lowest FDR values was performed using bar charts and bubble plots (Fig. 4).

**Fig. 4.**
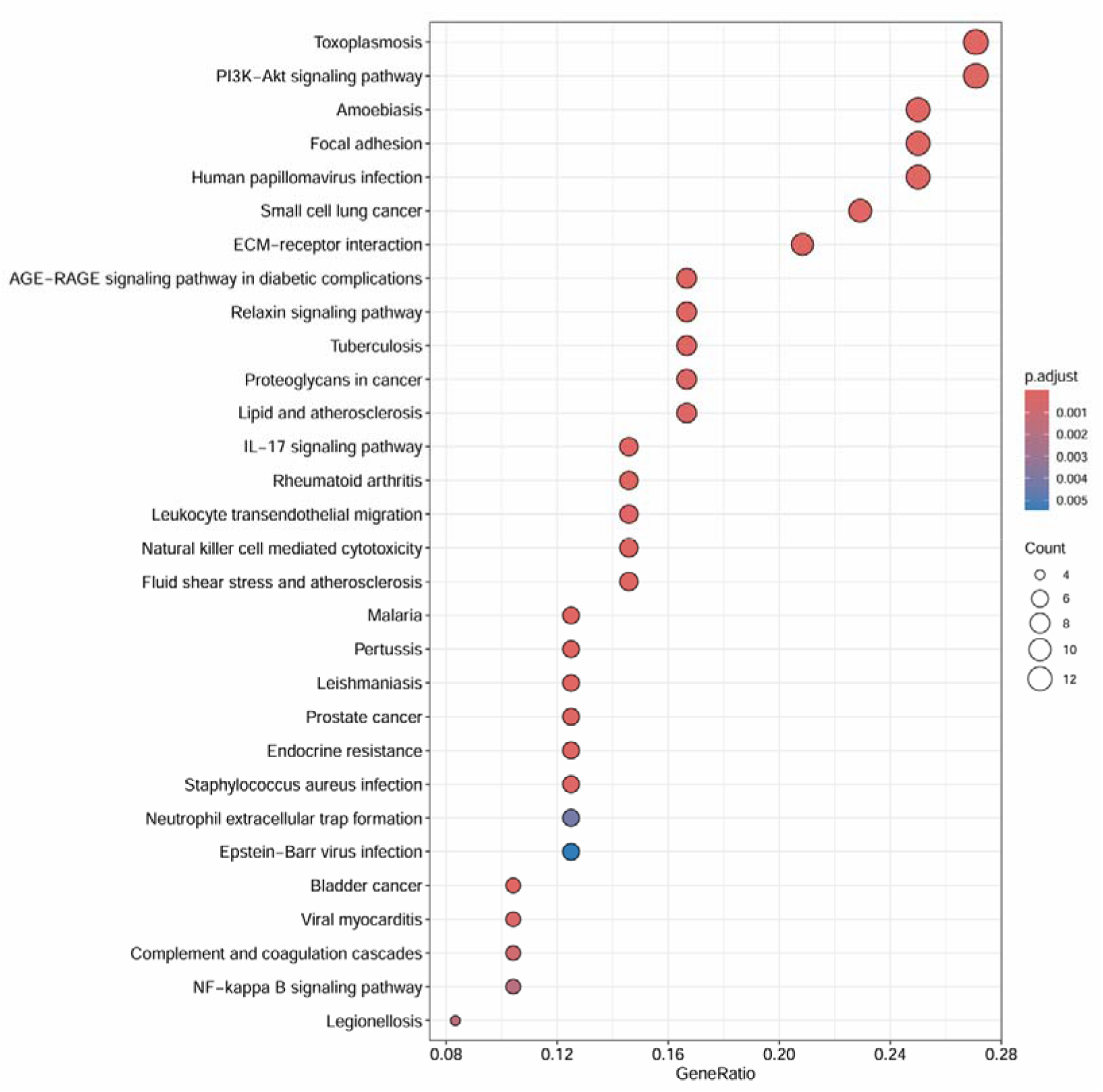
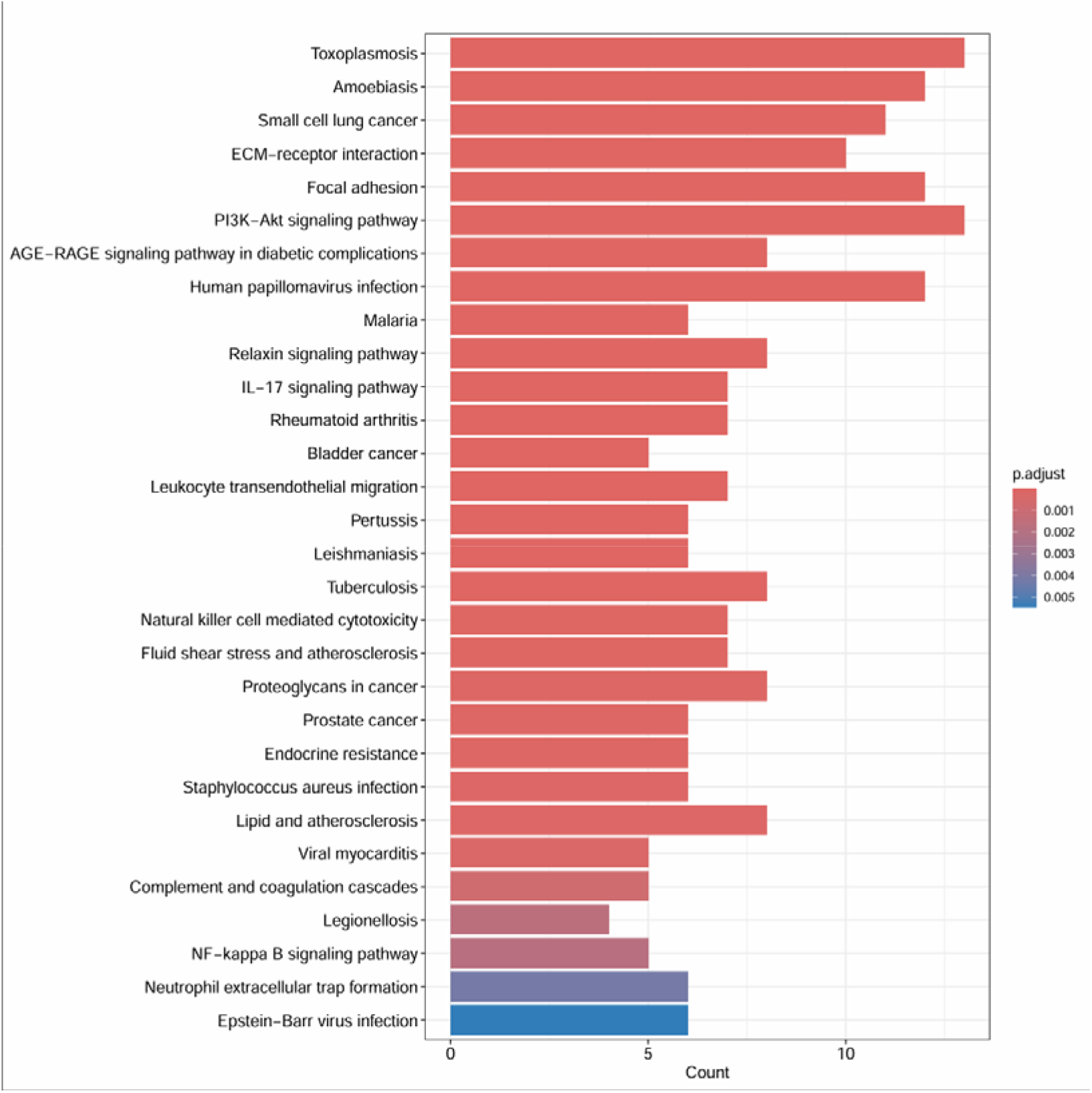
KEGG pathway enrichment analysis for genes intersecting between TBBPA and BP, performed using R. Significant enrichment was observed in pathways related to immune regulation and cellular stress, suggesting that TBBPA may contribute to BP pathogenesis through these regulatory pathways.

#### 3.3.2 Pathway Enrichment Analysis of Core Targets

Based on GO functional annotation and KEGG pathway enrichment results, biological mechanisms underlying TBBPA-induced BP were further analyzed. Significant enrichment in pathways associated with inflammation and immune responses, such as leukocyte migration, inflammatory response regulation, leukocyte transendothelial migration, IL-17 signaling pathway, and natural killer cell-mediated cytotoxicity, suggests these pathways mediate core biological events in TBBPA-induced BP. Additionally, enrichment in pathogen infection pathways, including toxoplasmosis, *Mycobacterium tuberculosis* infection, human papillomavirus infection, and Epstein-Barr virus infection, suggests immune dysregulation induced by chronic infection may trigger TBBPA-induced BP. Furthermore, significant enrichment in cancer-related pathways (small cell lung cancer, bladder cancer, prostate cancer, proteoglycans in cancer) indicates involvement of immune regulation and extracellular matrix remodeling mechanisms related to the tumor microenvironment. Lastly, enrichment in core pathways such as the PI3K-Akt signaling pathway, focal adhesion, ECM-receptor interaction, and NF-κB signaling pathway suggests these signaling disruptions are closely related to matrix homeostasis imbalance and abnormal inflammatory responses in TBBPA-induced BP.

### 3.4. Molecular Docking of TBBPA with BP Core Target Proteins

Molecular docking indicated that TBBPA exhibited strong binding affinity with all five core targets (TNF, CXCL8, MMP9, ICAM1, and ITGB1). Docking complexes exhibited low binding energies, with values ≤ -5 kcal/mol for TNF, CXCL8, MMP9, and ITGB1, indicating stable interactions. Analysis of 2D interactions showed TBBPA primarily engaged in van der Waals interactions with amino acid residues (tryptophan, phenylalanine, proline, valine, leucine) near active pockets, highlighting the importance of hydrophobic interactions in binding stability. Additionally, TBBPA formed specific π-π stacking interactions within the active pockets of CXCL8 and ITGB1. The prevalence of aromatic residues (phenylalanine, tryptophan) at binding sites suggests potential effects of TBBPA on fluorescence quenching. These findings visually demonstrate the tight binding between TBBPA and core BP proteins, providing a structural basis for its potential regulatory effects (Fig. 5).

**Fig. 5.**
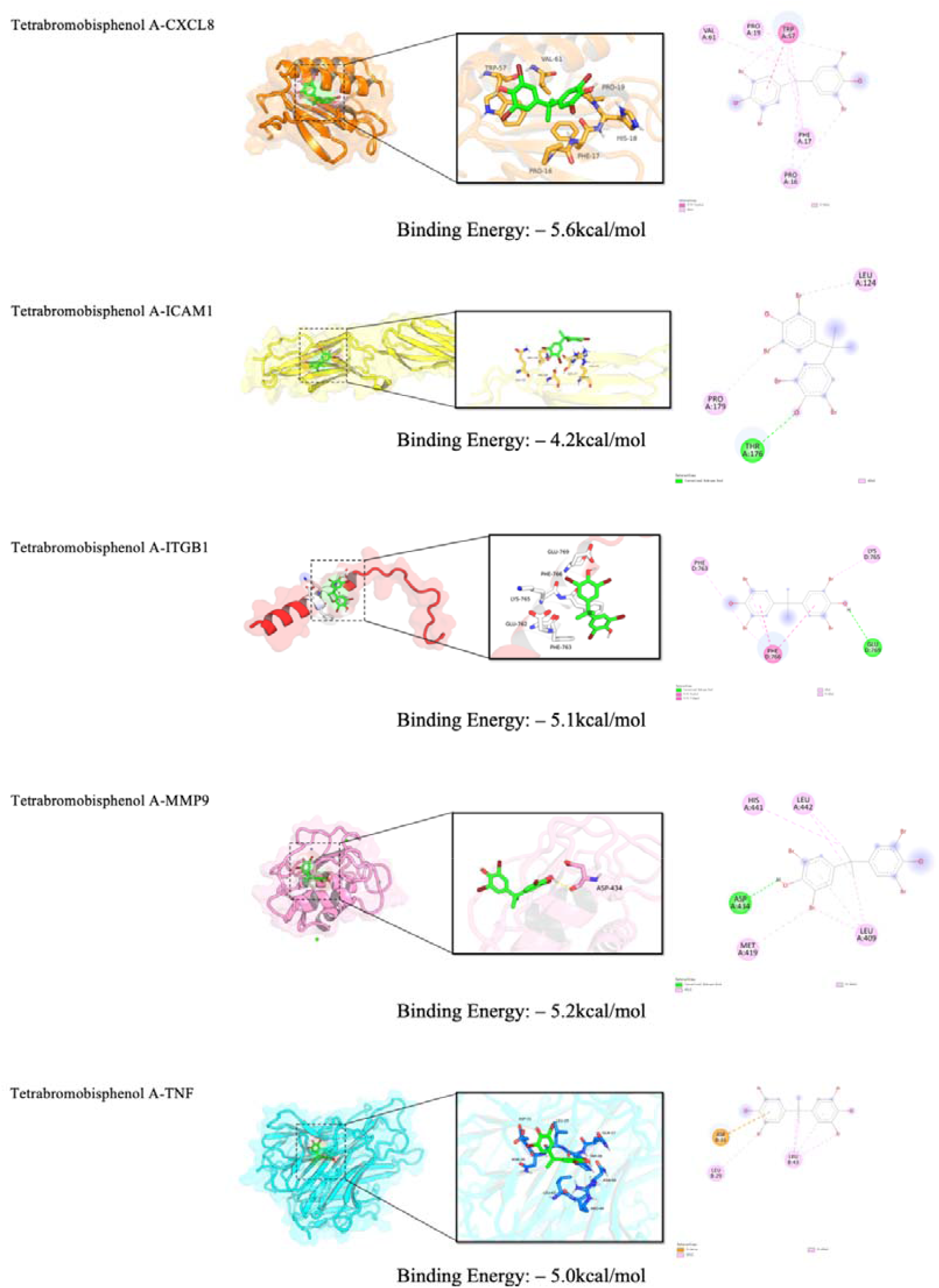
Visualisation of molecular docking.

### 3.5. MD Simulation of Core Targets

To validate the binding affinity of TBBPA with core targets, MD simulations were conducted (Fig. 6). Analysis indicated that TBBPA formed stable complexes with all three target proteins.

**Fig. 6.**
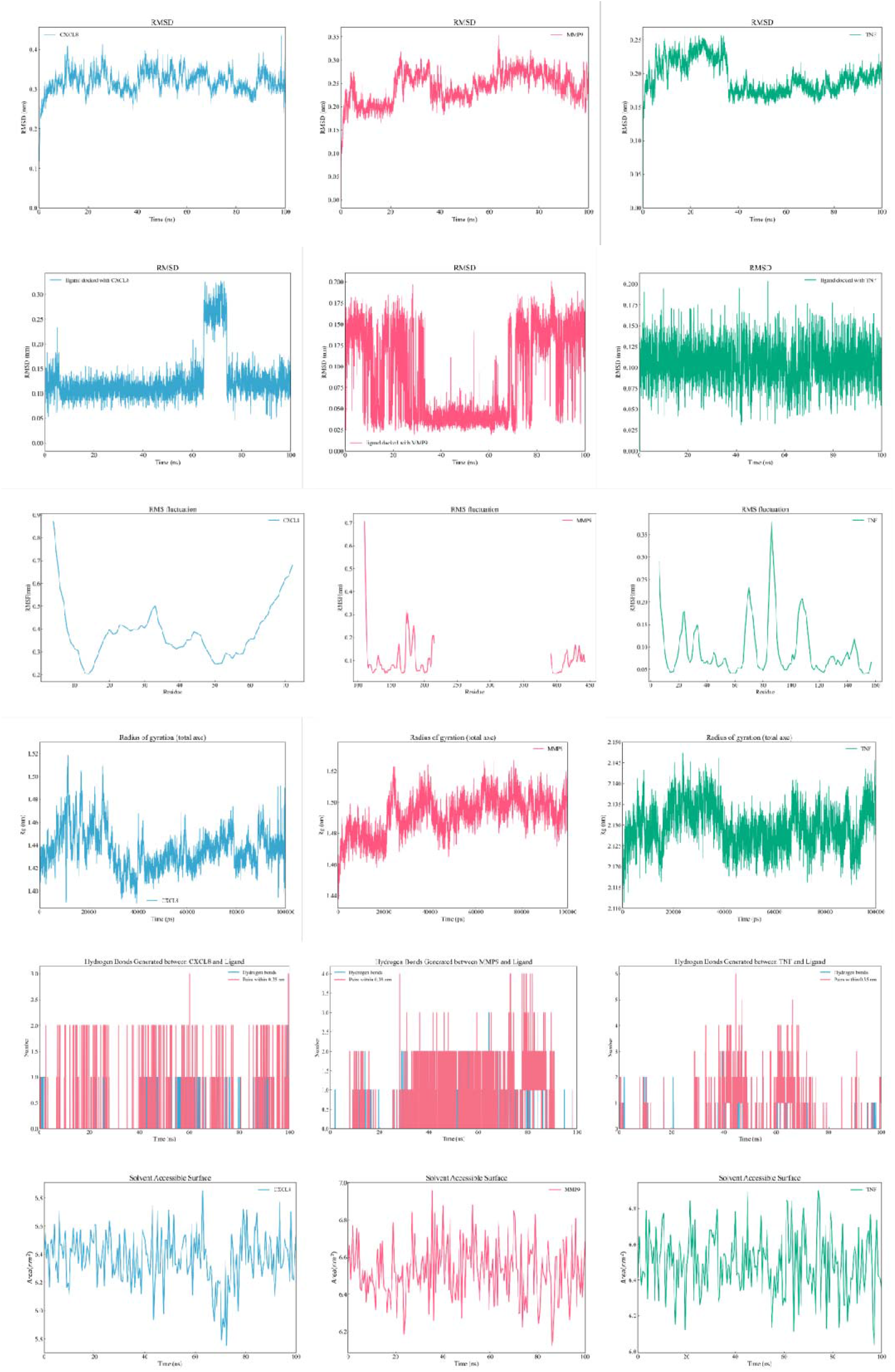
MD simulations.

The overall conformational stability was assessed using the RMSD. RMSD values for the protein backbones of the three protein-TBBPA complexes reached equilibrium plateaus after approximately 60 ns, 85 ns, and 40 ns, respectively. They then fluctuated stably around 0.30 nm, 0.25 nm, and 0.17 nm, indicating conformational stability after equilibration [20]. Ligand RMSD analysis showed minimal conformational changes during simulations, suggesting stable positioning within the binding pockets. Only the CXCL8 complex exhibited a minor fluctuation (ΔRMSD < 0.2 nm) at 65 ns, without conformational flipping.

Protein residue flexibility was analyzed using the RMSF. The N-terminal and C-terminal residues of all proteins exhibited higher fluctuations due to terminal flexibility. Significantly lower RMSF values were observed near residue 50 of CXCL8, residues 110-160, 195-200, and 400-405 of MMP9, and residues 40-60, 120-140, and 150-157 of TNF compared to other regions [21]. These low-fluctuation regions corresponded to ligand-binding pockets, suggesting ligand binding reduces conformational flexibility of these residues.

The overall compactness of the protein structures was evaluated using the radius of gyration (Rg). The Rg values of all three proteins remained stable throughout the 100 ns simulation, with fluctuations of less than 0.2 nm, indicating maintained structural integrity and compactness.

The stability of intermolecular interactions was assessed through hydrogen bond analysis. Throughout simulations, TBBPA formed 1-2 stable hydrogen bonds with CXCL8 and MMP9, and intermittently formed 1-3 hydrogen bonds with TNF. Additionally, stable hydrophobic and other non-covalent interactions were observed, highlighting their importance in maintaining complex stability.

Finally, the SASA of the ligand was analyzed to evaluate exposure within binding pockets. The SASA of TBBPA in all systems remained stable, without significant increases, indicating effective encapsulation within protein binding pockets. Notably, in the CXCL8 complex, ligand SASA transiently decreased during 65-73 ns, corresponding to minor RMSD fluctuations. This may indicate local conformational adjustments within the binding pocket, enhancing ligand binding.

## 4. Discussion

This integrated computational study demonstrates that TBBPA can target core proteins (TNF, CXCL8, MMP9, ICAM1, and ITGB1), potentially interfering with multiple inflammatory and immune-related pathways and thereby contributing to the pathogenesis of BP. This study constructs a systematic molecular framework linking TBBPA exposure to BP susceptibility.

Tumor necrosis factor (TNF) was identified as the top-ranked core target, a finding with significant biological relevance, as previous studies confirmed its key role in the inflammatory response of BP [22,23]. Similarly, chemokine CXCL8 and matrix metalloproteinase 9 (MMP9) are central to neutrophil-mediated blister formation [24,25]. The prediction that TBBPA can stably bind these proteins suggests it may modulate their activity, thereby influencing immune cell recruitment and tissue damage.

This study has limitations. Findings are derived from computational simulations and lack experimental validation. Subsequent studies should conduct binding assays, cell-based experiments, and *in vivo* studies to verify the functional effects of TBBPA and clarify its causal relationship with BP pathogenesis.

## 5. Conclusion

This study systematically investigated the potential molecular mechanisms by which TBBPA, a common brominated flame retardant, induces BP by integrating network toxicology, molecular docking, and MD simulations.

Using network toxicology, we identified 48 potential common targets linking TBBPA exposure to BP pathogenesis. PPI network analysis identified five core regulatory targets: TNF, CXCL8, MMP9, ICAM1, and ITGB1. These targets hold hub positions in the network and are closely linked to BP-related pathological processes, including inflammation, immune cell infiltration, and basement membrane disruption.

Functional enrichment analysis revealed that these targets significantly enrich immune-inflammatory pathways, such as leukocyte migration, inflammatory response regulation, and the IL-17 signaling pathway, along with infection- and cancer-related pathways. This suggests that TBBPA might reduce BP development thresholds by perturbing immune homeostasis and exacerbating chronic inflammation.

Molecular docking results indicated that TBBPA could stably bind all five core targets (TNF, CXCL8, MMP9, ICAM1, ITGB1), with binding energies ≤ -5 kcal/mol. The interactions were predominantly driven by hydrophobic interactions and specific π-π stacking. Subsequent MD simulations confirmed that complexes formed by TBBPA with TNF, CXCL8, and MMP9 remained stable during a 100 ns simulation. Protein structures remained compact, and the ligand was effectively encapsulated within binding pockets through stable hydrogen bonds and hydrophobic interactions.

In summary, this computational study proposes a systematic molecular framework indicating that TBBPA might contribute to BP pathogenesis in susceptible individuals. By targeting core proteins (TNF, CXCL8, and MMP9) and disrupting key signaling pathways related to immune inflammation, cell migration, and tissue remodeling, this study provides a novel mechanistic hypothesis and potential therapeutic targets. This advances our understanding of how environmental chemical pollutants influence autoimmune blistering diseases.

## CRediT authorship contribution statement

**Sun Kuo**: Conceptualization, Methodology, Software, Formal analysis, Investigation, Writing – original draft, Visualization. **Liu Yunxin**: Methodology, Validation, Software, Formal analysis, Investigation, Writing – review & editing, Visualization. **Zhao Hanqing**: Resources, Writing – review & editing, Supervision, Project administration.

## Data availability

The data supporting this study are available from public databases as cited in the manuscript.

## References

[1] X. Yang, P. Wei, Z. Wang, 2025. Recent advances in the genetics and innate immune cells of bullous pemphigoid. Front. Immunol. 16, 1530407. 10.3389/fimmu.2025.1530407.

[2] K. Liu, J. Li, S. Yan, W. Zhang, Y. Li, D. Han, A review of status of tetrabromobisphenol A (TBBPA) in China, Chemosphere. 148 (2016) 8–20. 10.1016/j.chemosphere.2016.01.023.

[3] A.L. Hopkins, Network pharmacology: the next paradigm in drug discovery, Nat. Chem. Biol. 4 (2008) 682–690. 10.1038/nchembio.118.

[4] X.Y. Meng, H.X. Zhang, M. Mezei, M. Cui, Molecular docking: a powerful approach for structure-based drug discovery, Curr. Comput. Aided Drug Des. 7 (2011) 146–157. 10.2174/157340911795677602.

[5] T. Cui, Y. Zhou, T. Wang, 2025. Recent advances in artificial intelligence-driven biomolecular dynamics simulations based on machine learning force fields. Curr. Opin. Struct. Biol. 95, 103191. 10.1016/j.sbi.2025.103191.

[6] B. Zdrazil, E. Felix, F. Hunter, E.J. Manners, J. Blackshaw, S. Corbett, M. de Veij, H. Ioannidis, D.M. Lopez, J.F. Mosquera, M.P. Magarinos, N. Bosc, R. Arcila, T. Kizilören, A. Gaulton, A.P. Bento, M.F. Adaasme, P. Monecke, G.A. Landrum, A.R. Leach, The ChEMBL Database in 2023: a drug discovery platform spanning multiple bioactivity data types and time periods, Nucleic Acids Res. 52 (2024) D1180–D1192. 10.1093/nar/gkad1004.

[7] Z.J. Yao, J. Dong, Y.J. Che, M.F. Zhu, M. Wen, N.N. Wang, S. Wang, A.P. Lu, D.S. Cao, TargetNet: a web service for predicting potential drug-target interaction profiling via multi-target SAR models, J. Comput. Aided Mol. Des. 30 (2016) 413–424. 10.1007/s10822-016-9915-2.

[8] A. Daina, O. Michielin, V. Zoete, SwissTargetPrediction: updated data and new features for efficient prediction of protein targets of small molecules, Nucleic Acids Res. 47 (2019) W357–W364. 10.1093/nar/gkz382.

[9] UniProt Consortium, UniProt: the Universal Protein Knowledgebase in 2025, Nucleic Acids Res. 53 (2025) D609–D617. 10.1093/nar/gkae1010.

[10] G. Stelzer, N. Rosen, I. Plaschkes, S. Zimmerman, M. Twik, S. Fishilevich, T.I. Stein, R. Nudel, I. Lieder, Y. Mazor, S. Kaplan, D. Dahary, D. Warshawsky, Y. Guan-Golan, A. Kohn, N. Rappaport, M. Safran, D. Lancet, The GeneCards Suite: From Gene Data Mining to Disease Genome Sequence Analyses, Curr. Protoc. Bioinform. 54 (2016) 1.30.1-1.30.33. 10.1002/cpbi.5.

[11] J.S. Amberger, C.A. Bocchini, F. Schiettecatte, A.F. Scott, A. Hamosh, OMIM.org: Online Mendelian Inheritance in Man (OMIM®), an online catalog of human genes and genetic disorders, Nucleic Acids Res. 43 (2015) D789–D798. 10.1093/nar/gku1205.

[12] Y. Zhang, Y. Zhou, H. Xu, W. Jiang, B. Li, D. Lai, C. Wan, S. Wang, M. Zhao, Y. Tan, S. Lu, T. Fan, X. Liu, F, Zhu, Y. Chen, Therapeutic target database 2026: facilitating targeted therapies and precision medicine, Nucleic Acids Res. 54 (2026) D1692–D1701. 10.1093/nar/gkaf1154.

[13] D. Otasek, J.H. Morris, J. Bouças, A.R. Pico, B. Demchak, 2019. Cytoscape Automation: empowering workflow-based network analysis. Genome Biol. 20, 185. 10.1186/s13059-019-1758-4.

[14] G. Yu, L.G. Wang, Y. Han, Q.Y. He, clusterProfiler: an R package for comparing biological themes among gene clusters, OMICS. 16 (2012) 284–287. 10.1089/omi.2011.0118.

[15] M. Carlson, org.Hs.eg.db: Genome wide annotation for Human. R package version 3.15.0. Bioconductor (2022).

[16] M. Kanehisa, M. Furumichi, Y. Sato, M. Ishiguro-Watanabe, M. Tanabe, KEGG: integrating viruses and cellular organisms, Nucleic Acids Res. 49 (2021) D545–D551. 10.1093/nar/gkaa970.

[17] H. Wickham, ggplot2: Elegant Graphics for Data Analysis, Springer-Verlag, New York, 2016.

[18] A. Kassambara, ggpubr: ‘ggplot2’ Based Publication Ready Plots. R package version 0.5.0 (2022).

[19] A. Sinelnikova, D.V. Spoel, NMR refinement and peptide folding using the GROMACS software, J. Biomol. NMR. 75 (2021) 143–149. 10.26434/chemrxiv.13637819.

[20] E.A. Coutsias, M.J. Wester, RMSD and Symmetry, J. Comput. Chem. 40 (2019) 1496–1508. 10.1002/jcc.25802.

[21] C.S. Sharanya, D.S. Wilbee, S.N. Sathi, K. Natarajan, 2024. Computational screening combined with well-tempered metadynamics simulations identifies potential TMPRSS2 inhibitors. Sci. Rep. 14, 16197. 10.1038/s41598-024-65296-7.

[22] C. Feliciani, P. Toto, P. Amerio, S.M. Pour, G. Coscione, G. Shivji, B. Wang, D.N. Sauder, In vitro and in vivo expression of interleukin-1alpha and tumor necrosis factor-alpha mRNA in pemphigus vulgaris, J. Invest. Dermatol. 114 (2000) 71–77. 10.1046/j.1523-1747.2000.00835.x.

[23] F. Ameglio, L. D’Auria, P. Cordiali-Fei, A. Mussi, L. Valenzano, G. D’Agosto, C. Ferraro, C. Bonifati, B. Giacalone, Bullous pemphigoid and pemphigus vulgaris: correlated behaviour of serum VEGF, sE-selectin and TNF-alpha levels, J. Biol. Regul. Homeost. Agents. 11 (1997) 148–153. 10.1038/sj.ijo.0800496.

[24] E. Schmidt, S. Reimer, S. Jainta, E.-B. Bröcker, D. Zillikens, N. Kruse, M.P. Marinkovich, G.J. Giudice, Autoantibodies to BP180 associated with bullous pemphigoid release interleukin-6 and interleukin-8 from cultured human keratinocytes, J. Invest. Dermatol. 115 (2000) 842–848. 10.1046/j.1523-1747.2000.00141.x.

[25] N. Cirillo, S.S. Prime, 2021. A Scoping Review of the Role of Metalloproteinases in the Pathogenesis of Autoimmune Pemphigus and Pemphigoid, Biomolecules. 11, 1506. 10.3390/biom11101506.

